# Fast and lightweight cell atlas approximations across organs and organisms

**DOI:** 10.1101/2024.01.03.573994

**Authors:** Ying Xu, Joanna Ahn, Fabio Zanini

## Abstract

Omic technologies at single-cell resolution are reshaping our understanding of cellular diversity. The generation of cell atlases that capture the cellular composition of an entire individual is progressing rapidly. However, the science of organising and extracting information from these atlases is still in its infancy and for many biomedical researchers atlas exploration remains challenging. Here, we leveraged extensive experience in single-cell data analytics to pinpoint three major accessibility barriers to cell atlases, related to (i) programming skill or language, (ii) scalability, and (iii) dissemination standards. To help researchers overcome these barriers, we developed cell atlas approximations, a computational approach enabling the analysis of cell atlases across organs and organisms without programming skills, rapidly, and at scale. The web interface at https://atlasapprox.org facilitates the exploration of cell atlases in 19 species across the tree of life through a chatbot driven by frontend natural language processing. In parallel, application programming interfaces streamline data access for computational researchers and include specialised packages for Python, R, JavaScript, and Bash. Supported queries include marker gene identification, cross-organ comparisons, cell embeddings, gene sequences, searches for similar features, and bidirectional zoom between cell types and cell states. Most queries are answered in less than 1.5 seconds thanks to lossy data compression algorithms based on cell annotations and similarity graphs. Compared to traditional cell atlas analysis, this approach can reduce data size by more than 100 times and accelerate workflows by up to 100,000 times. Atlas approximations aim to make the exploration of cell atlases accessible to anyone in the world.

## Introduction

Single-cell omic technologies such as scRNA-Seq and scATAC-Seq can measure cell heterogeneity with both granularity and scalability [1]. In parallel with efforts on individual diseases and developmental stages, a growing number of studies are collecting reference data covering the entire body of multicellular organisms across the tree of life [2–12]. These whole-body cell atlases constitute an incredibly rich source of biological information about the types, subtypes, and states of living cells, connecting genotype with molecular phenotype (e.g. gene expression). While atlas data generation is progressing in both animals and plants, research on how to best summarise, explore, and extract information from atlas data is in early stages.

Access to cell atlas data is currently hampered by significant barriers. Many reference atlases are massive in size and complexity, making them slow to load, query, and interpret. Furthermore, many exploratory analyses on cell atlases require advanced scientific programming skills, thereby excluding much of the biomedical community. Both elements discriminate against researchers in low-income or low-skill settings, for whom acquiring suitable computer hardware and training data scientists can be challenging. The issue is also exacerbated by a lack of standardised online databases and file formats, unclear data preprocessing and, sometimes, inaccurate cell type annotation. As a result, many potential discoveries remain hidden from the scientific community, thwarting important progress in both basic cell biology and molecular medicine.

Several tools and websites have been created to simplify the exploration of single cell omic data, including ShinyCell [14], cellxgene [15], and SPEED [16]. These programs share the following design pillars: (i) An emphasis on cell embeddings as the primary user interface (UI) to atlas data [13]; (ii) User interactions based on HTML buttons, including dropdown menus, “submit” buttons, and buttons to colour the embedding in various ways; (iii) A “no cell left behind” approach which aims to visualise each and every individual cell of the atlas. At the same time, support for biologically relevant functionality such as marker genes, access to gene sequences, and comparison across tissues is patchy at best. Notably, none of these tools were designed specifically for whole-body cell atlases.

Existing web portals do not generally offer an easy way to access or download the data being visualised, which degrades user experience. As soon as a question exceeds the expressiveness of HTML buttons, the user’s only option is to locate the dataset on an online database, download Gigabytes of data, and start a separate secondary analysis from scratch. Cellxgene Census [15], a set of Python and R APIs for cellxgene, has recently shown promising results, however it is heavily focused on human and disease-related data, and the functionality is quite limited. Moreover, Census still needs to stream data from thousands or millions of cells across the network, which can take minutes to days depending on the user’s internet connection. Of course, reliance on good internet bandwidth also biases against researchers in low-resource settings.

Here, we present atlas approximations, highly compressed representations of whole-body cell atlases aimed at extracting the essential bits of information from each atlas and disseminating them to both machines and humans independently of user background, location, hardware quality, and computational literacy. Our database includes 19 species, contains data on both gene expression and chromatin accessibility, and offers functionality not available elsewhere. Instead of HTML buttons, a chatbot was built to interpret questions by humans and provide an answer, in text and graphic form, within 1.5 seconds. Application programming interfaces (APIs) were also implemented to respond to queries from any programming language within 1 second, with dedicated packages in Python, R, Bash, and JavaScript. Two approximation algorithms were designed to enable data exploration at different levels of resolution. All components (data, algorithms, machine interface, chatbot, and web interface) are open source, thoroughly tested, and designed for easy recycling.

## Results

### Identifying the accessibility barriers to cell atlases

We have accrued years of experience analysing single cell omic data across multiple species and diverse contexts including infectious disease [17–19], development [20,21], cancer [22], ageing [23], and neonatal disease [21]. In each of those studies, comparison of gene expression or chromatin accessibility data with a published cell atlas such as Tabula Sapiens [3] was not easy to perform. We realised that only a small fraction of biomedical researchers would be able to explore a cell atlas at all because of three major accessibility barriers.

#### Barrier 1 (Programming skill or language)

Atlas explorations require coding skills. This barrier exists independently of high-quality documentation for popular single cell analytical software such as scanpy [24] and seurat [25]. Once a researcher is familiar with Python syntax and “the stack” based on numpy [26], scipy [27], matplotlib [28], pandas [29], igraph [30], and Jupyter [31], learning scanpy is straightforward. Nonetheless, the vast majority of biomedical researchers are not familiar with these tools nor their equivalents in other programming languages (**Figure 1A**).

**Figure 1.**
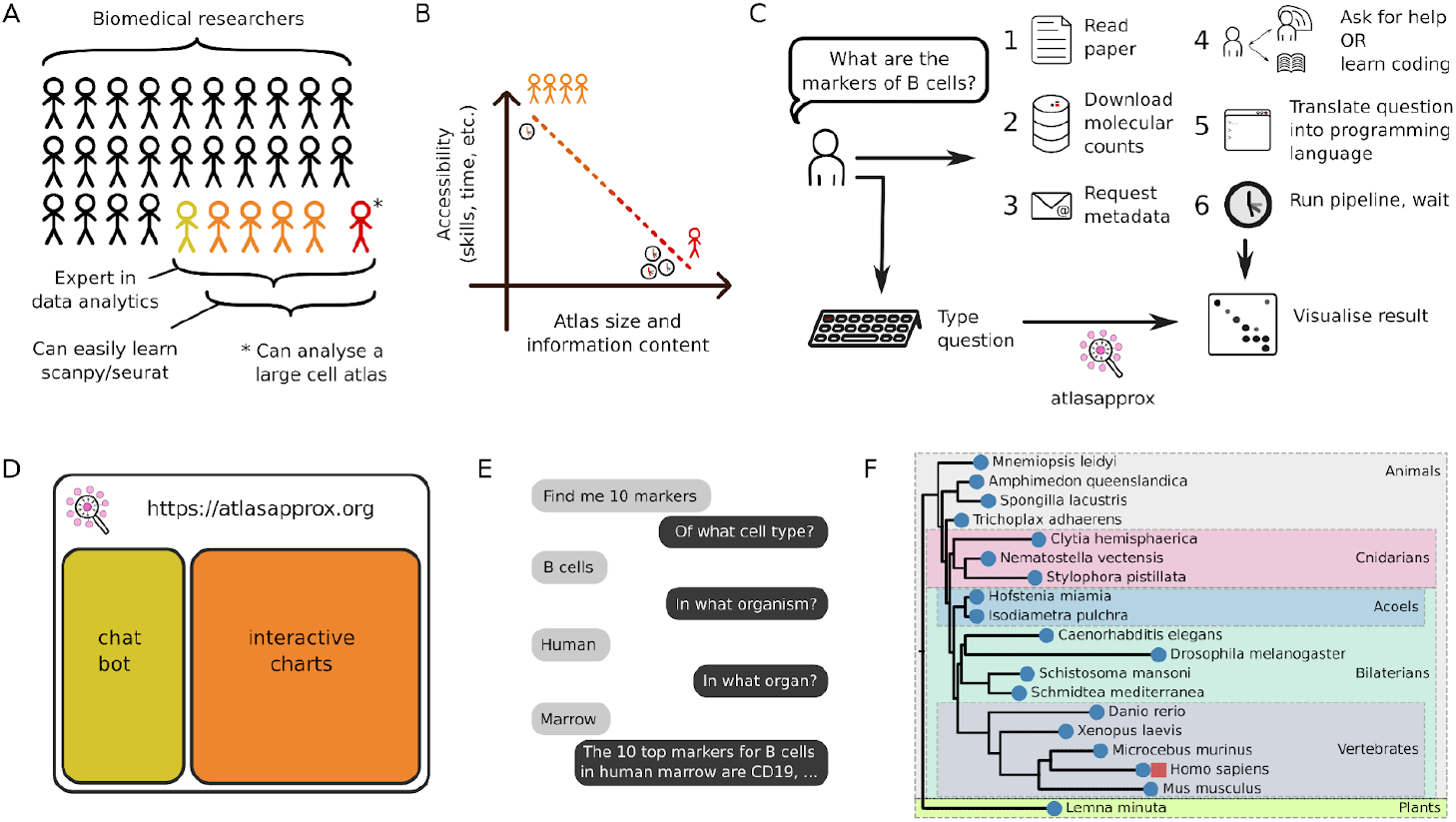
Approximations aim to overcome the three major accessibility barriers to cell atlases. (A) Because browsing most cell atlases requires scalable scientific programming, only a small fraction of the biomedical community can actually access them. (B) Increasing atlas size and information content reduces the accessibility including in terms of skills and time requirements. (C) Typical traditional workflow to access and analyse cell atlas data. (D) Schematic of the web interface of cell atlas approximations. (E) Typical conversation with the atlasapprox chat bot. (F) Species with currently available cell atlas approximations for scRNA-Seq (blue circles) and scATAC-Seq (red square) across the tree of life. Branch lengths are proportional to the number of intermediates in Open Tree of Life [41]. A few clades are indicated by coloured boxes and labelled on the right.

#### Barrier 2 (Scalability paradox)

The larger and more informative the atlas, the lower its accessibility, because larger atlases, sometimes incorporating dozens of tissues or spanning multiple measurement modalities, are harder to download, store, and explore. They require more powerful hardware and especially optimised software or take longer to analyse. Yet, they also contain the most potential discoveries, a paradox that has not been solved to date (**Figure 1B**).

#### Barrier 3 (Poor dissemination standards)

In our experience, the typical journey to explore a published atlas is slow and frustrating (**Figure 1C**). It starts with finding a publication where the data is described, which should contain a link to a table of counts (cells x features) hosted on one of many, mutually idiosyncratic databases. Key metadata (e.g. cell annotations, embedding coordinates) often have to be requested via email, which takes weeks to months and can be unsuccessful. Subsequently, a custom software pipeline needs to be written to standardise or correct both counts and cell type annotations. Afterwards, to extract simple information from each atlas, human questions must be translated into machine language (see barrier 1). In addition to the above, because of the massive size of some atlases, it could take half an hour to answer a question as simple as comparing one single cell type across multiple tissues (see barrier 2).

### Designing an accessible interface to whole-body cell atlases

Ideally, a user with no coding skills would type a biological question in an online prompt (e.g. “What are 10 marker genes of rod cells in the frog eye?”) and get an answer from an arbitrarily large cell atlas within seconds (**Figure 1C**). Cell atlas approximations were designed to embody this ideal and overcome all three accessibility barriers at once. Although the software stack behind atlas approximations is complex (**Supplementary Figure 1**), they are used by most researchers via the web interface, freely available at https://atlasapprox.org. The interface is vertically split in two: a text area on the left and a plotting area on the right (**Figure 1D**). The plotting area is used for interactive data visualisations. The text area hosts conversations with an “intelligent” chatbot, who is in charge of understanding the researcher’s question in English, translating it into machine code, and providing an answer from the atlas data within 1.5 seconds. A single query typically involves 1-5 exchanges (**Figure 1E**). The data used in atlas approximations currently covers 19 organisms across the tree of life (**Figure 1F**), with more scheduled for the future.

### Natural language processing on the frontend enables equitable accessibility

The chatbot is a key component of atlas approximations because it overcomes the first accessibility barrier, i.e. it opens atlas explorations to non-programmers. It has two remarkable features.

First, the bot uses natural language processing (NLP), a branch of artificial intelligence (AI), to understand the user’s general intent and to recognise any named entities such as the names of organisms, organs, cell types, genes, and so on (**Figure 2A**). If the intent is clear but not all entities have been specified in the user query, the bot seeks to fill the missing “information slots” via follow-up questions (**Figure 1E**). Once a complete query is negotiated, a machine action is triggered, usually involving requests to a separate server hosting the atlas data. The data is then formatted into an answer, which includes both text (left side of the page) and graphics (right side).

**Figure 2.**
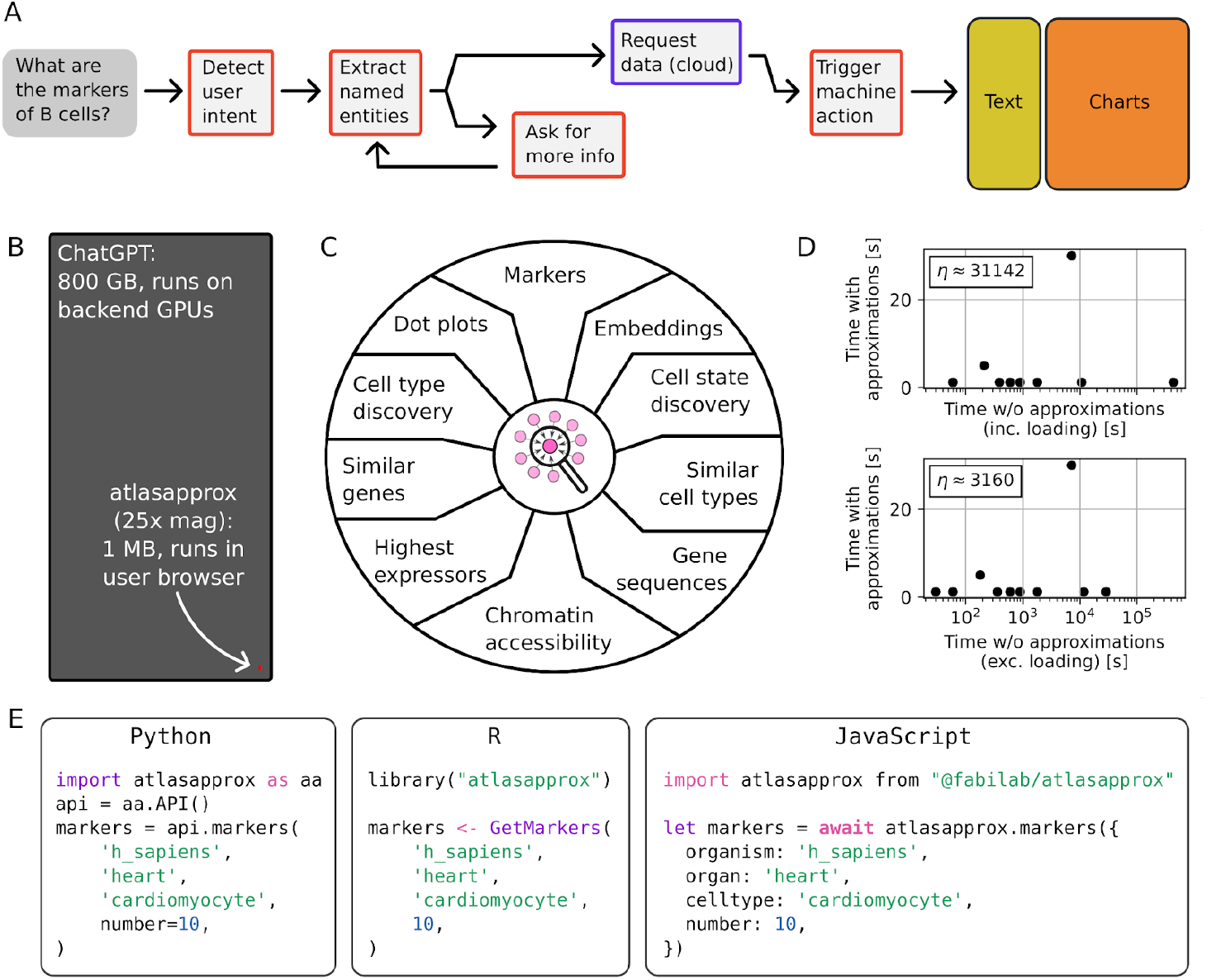
Chatbot and application programming interfaces to cell atlas approximations. (A) Flow diagram of the logic behind a single request as it travels through the chatbot. (B) Visualisation of the size of the ChatGPT [32] versus the atlas approximations chatbot. (C) Some of the intents understood by the chatbot. (D) Anecdotal runtimes for a range of queries using atlas approximations versus full atlases. (E) Example listings of the atlas approximations application programming interfaces (APIs) in three programming languages.

Second, the bot is designed differently from the popular NLP bot ChatGPT [32]. ChatGPT ingests the user’s utterances and transmits them to a remote server, which runs an 800 GB model using arrays of GPUs. The model’s owner can thereby harvest both subscription fees and data from their own users. In contrast, our bot is around 1 MB in size (**Figure 2B**) and runs entirely within the user’s browser (no GPU needed). No data is sent to any remote server with the exception of the machine queries to the atlas data server. This architecture ensures sustainable and equitable access for researchers from low-income settings, because it does not rely on a subscription nor accumulates user utterance data. In the same spirit, the full web interface is a zero-backend static page with zero hosting cost: avid readers are welcome to fork it and adapt it to their own needs.

The chatbot recognises a spectrum of biological intents including marker gene identification, cross-organ comparisons, dot plots for cell types, and embeddings for cell states (**Figure 2C**). The user can also start from one gene of interest and search for additional genes with similar patterns of expression across cell types and tissues. Also, chromatin accessibility is available for human data already and planned for additional species. Gene sequences can also be retrieved and downloaded. A unique functionality is the ability to transition between two levels of granularity, starting at the cell type level and then zooming into cell states, and vice versa.

Anecdotal runtimes for typical operations related to those intents indicated speed-ups between 10 and 100,000 times compared to traditional atlas analysis (**Figure 2D**). All of these intents could be answered much faster via atlas approximations than by locating and browsing the atlas using traditional methods, however the difference was particularly stark for intents that involved either comparisons across multiple organs or massive atlases, as expected by the scalability barrier (**Figure 1B**). The only case where atlas approximations took longer than a couple of seconds was related to requests that require multiple steps. Importantly, that limitation can be mitigated by engineering additional intents for the chatbot.

### Application programming interfaces standardise machine access

While the chatbot attends to human language queries, atlas approximations also provide application programming interfaces (APIs) to simplify machine queries to cell atlas data. A REST API was constructed to ensure accessibility from any programming language. Dedicated interfaces in Python, R, JavaScript, and Bash were also built to improve accessibility for users of those languages. API requests take only a handful of lines of code in any language (**Figure 2E**). Responses typically arrive within 900 ms of wall time.

### Lossy compression enables fast and scalable atlas queries

How can we provide answers within seconds from atlases with half a million cells? The answer is aggressive lossy compression. For most researchers, cells or nuclei as sampled in a specific experiment are not interesting *per se* but because they act as concrete realisations of an abstract cell type or state. Although the scientific community has not reached an agreement about the exact meaning of “cell type”, on a coarse level the concept is intuitive: neurons are much unlike fibroblasts in appearance, gene expression, chromatin accessibility, and function. Vice versa, on a very granular level, if the transcriptomes from two cells are 99% identical, data size can be halved by averaging them while retaining *most biological* information. Smaller data is, of course, easier and faster to download and manipulate. Cell atlas approximations leverage these ideas via two lossy compression algorithms (**Figure 3A**).

**Figure 3:**
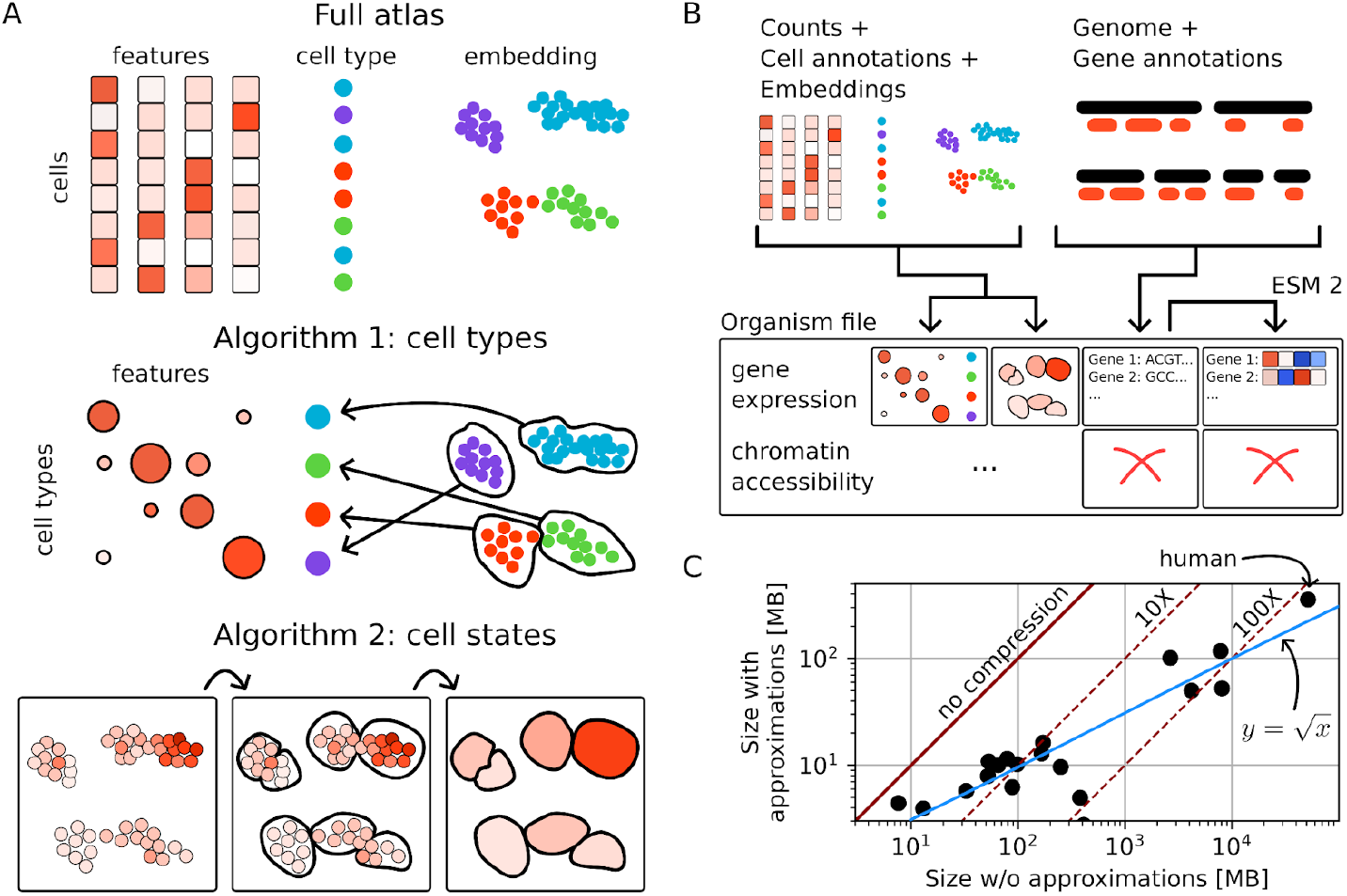
Design and performance of atlas approximations’ lossy compression algorithms. (A) Schematic of full atlas data (top) and compression algorithms 1 (middle) and 2 (bottom). Input data include the cell x feature count matrix, standardised cell type annotations (only for algorithm 1), and embedding coordinates (only for algorithm 2). (B) Schematic of the file format used to store cell atlas approximations, which includes molecular counts from one or multiple modalities, cDNA or peptide sequences, and ESM 2 embedings of peptides [34]. (C) Compression performance of atlas approximations visualised as size of each file versus total size of the original data. The solid red line indicates no compression. The dashed lines indicate constant compressions at 10x and 100x. The solid blue line indicates the empirical line shown in the equation.

Compression algorithm 1 averages the molecular counts (e.g. gene expression) within each cell type (**Figure 3A, middle**). Cell type annotations were initially taken from each publication, however they were later modified according to several criteria. Gross misannotations were corrected (e.g. epithelial cells marked as endothelial). Annotations that were very specific were grouped into coarser categories, especially if their original name was uninformative (e.g. “fibroblast III” would become “fibroblast”). Annotations that were too generic to be useful were split into subcategories using known marker genes whenever possible (e.g. “immune” would be split into “B cells”, “T cells”, and so on). Cell types that should be physiologically restricted to one tissue but were somehow found across many organs were blacklisted from all organs except the native one (e.g. “sperm” would be blacklisted in, say, the brain). Annotation names that described anatomical location were simplified (e.g. “fibroblast of heart” would become “fibroblast”). Finally, spelling mistakes and capitalisation were fixed throughout. This algorithm was designed to provide a biology-based approximation of an atlas that is coarse enough to be both intuitive and very fast to access.

Compression algorithm 2 also averages the molecular counts, but this time within finer cell states. To define a cell state in this context, we used the cell embeddings from each publication (if unavailable, we generated new ones). We then performed a modified version of K-means clustering in the embedding space, aiming to get around 3-4 times more clusters than the number of cell types in that tissue. Clustering in embedding space had two main advantages. First, the resulting clusters were always convex, so convex hulls can be used as visually intuitive proxies for measurements within each cell state (**Figure 3A, bottom right**). Second, if two cells had very similar profiles, they were likely to be projected near each other by the embedding algorithm [13] and therefore to be merged by the compression algorithm. Algorithm 2, which does not use cell type annotation, was designed to provide more granularity than algorithm 1. The cost is biological interpretability: while coarse cell types used by algorithm 1 have biologically meaningful names, cell states used by algorithm 2 have no names and can contain cells from multiple types. We mitigate this compromise by providing the composition of each cell state in terms of coarse cell types.

### Disseminating atlas approximations

Full atlases are often disseminated as either a monolithic database-like file (e.g. h5ad, rds) that includes both counts and metadata, or as a collection of files in different formats (e.g. matrix market format for the counts, comma-separated values for metadata). Although atlas approximations are mainly accessed via the chatbot or APIs, we designed a file format to foster dissemination in specific settings (e.g. bulk access). We mimicked h5ad [24] and store counts and metadata for both compression algorithms in an HDF5 file[33] (**Figure 3B**). We also include full cDNA or peptide sequences for all genes used in each atlas, which helps with cross-organism comparisons, such as finding “sister” cell types between evolutionarily distant species. In addition to the sequences, we also include 1024-dimensional ESM2 peptide sequence embeddings [34], enabling rapid comparison of proteins across the entire tree of life. During file writing, all data is chunked and passed through a lossless level-22 zstdandard ultracompression [35], ensuring minimal file size and fast decompression times despite slow compression times. Chromatin accessibility data is further quantised using 8-bit unsigned integers with uneven binning. The overall size of atlas approximation files compared to full atlases follows an empirical law of 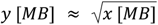 (**Figure 3C**). Because of this sublinear trend, the largest atlases - which have the highest accessibility barrier (**Figure 1B**) - are compressed the most. For instance, a whole human cell atlas including gene expression and chromatin accessibility from two dozen tissues plus the sequences of all human genes requires ∼400 MB versus ∼50 GB of the original data, all the while retaining *most biological* information (**Figure 3C**).

## Discussion

Whole-body cell atlases are being collected at an increasingly rapid pace, yet making such massive information vaults accessible to the biomedical community remains a formidable challenge. State-of-the-art approaches such as sparse matrices and disk-backed data loading [24] are useful but insufficient to address the fundamental accessibility barriers to cell atlases. These barriers are felt most acutely by biological domain experts, the very same people who would benefit the most from cell atlases’ information. Oncology and immunology are just two of many research areas that would be accelerated by a streamlined, code-free, and rapid interface to cell atlases. Cell atlas approximations were designed to bridge this accessibility gap.

The frontend chatbot was motivated by the language barrier and inspired by human-machine communication tools such as Amazon Alexa, Apple Siri, and Google Home. Unlike those commercial products, our chatbot does not transmit the user’s sentences to any remote server and is trained by a small corpus written by ourselves, thereby avoiding critical ethical issues with large language models [36]. In contrast to ChatGPT, our chatbot is designed to trigger machine actions rather than generate prose, which avoids AI hallucination [37]. Compared to other single-cell online portals, the chatbot avoids cluttering the screen with dozens of small buttons [15,16]. It is also equitable towards users from low-resource settings and flexible enough for future adaptations to languages beyond English.

The second and third accessibility barriers, i.e. lack of scalability and standards, prompted the design of the lossy compression algorithms and cross-language APIs. Compression ratios intuitively improve with growing atlas size because after most cell types have been sampled, deeper cell sampling yields diminishing returns in terms of novel biology [38]. Conversely, discarding nonessential information is likely to accelerate single-cell analytics by directing the researcher’s attention more effectively. This idea mirrors geographic atlases, which, understandably, do not depict every grain of sand. In the tree of life, each species is usually shown only once. A comparable scenario is commonly observed in online behaviour: when information becomes overly abundant, user attention gains importance instead.

The issue of cell type annotation is thorny because individual researchers have been unable to agree on a common definition of “cell type” and “cell state”. Therefore, many mathematical models including statistical approaches [39] and deep learning [40] use cell type annotations as *bona fide* training labels. Based on our effort reanalysing 20 whole-body cell atlases of all sizes and from diverse organisms, such faith is likely misplaced. Many issues were identified during our annotation processing, from uninformative names to gross negligence. In the light of that experience, models trained on uncorrected annotations from cell atlases are liable to major inference mistakes. In this study, we partially eschewed the problem by using coarse, intuitive annotations for compression algorithm 1 and no annotations for algorithm 2.

This study has several limitations. Cell atlas approximations, being lossy compressions, do not retain the full information from each individual cell. Cell type- or state-level information is sufficient for many purposes, but specific niches such as antigen-specific B or T cell immunology are not covered. In principle, more granular approximations beyond cell state could be conceived, ultimately interpolating down to individual cells. Furthermore, approximations are currently restricted to adult, healthy individuals as representatives for an entire species. Extensions to development and disease will be the topic of future work, as will be additional species beyond the current 19. Finally, the chatbot and data format developed for this study were custom tailored to the available atlases, therefore not generally applicable to each researcher’s own data. Efforts to overcome this constraint and enable natural language-driven exploration of custom single cell data sets are underway.

Recent history of cell biology seems to parallel the earlier canon of evolutionary biology. Initially, morphology was used to identify species and cell types, a methodology that was slow and sometimes inaccurate but accessible to everyone. Subsequently, sequencing technologies led researchers to multiple-sequence alignments and single-cell omics, which are more quantitative but less accessible to non-computational researchers. Finally, after the utility and limitations of the “species’’ concept became accepted, projects such as the Open Tree of Life [41] arose to provide accessible universal knowledge for the benefit of all. Similarly, as the advantages and limitations of the “cell type” concept are understood better each day, we hope that cell atlas approximations might become an equally useful tool towards a universal understanding of cell biology across the tree of life.

## Methods

### Atlas collection

Whole-body single cell atlases were discovered via online search engines and emails to academics in the single cell field. Single cell data sets that were specific to one tissue, disease, or developmental stage were not considered. Links to processed data (count matrices) were identified. If available, data and/or metadata were downloaded directly, otherwise email communication with the lead and/or senior authors of the publication was initiated. Whenever both data and metadata could be located, attempts were made at reconciling them and thereby including them into our list.

### Data preprocessing

Molecular count tables were reverse engineered to restore raw counts, undoing all downstream preprocessing done for the publication. Data was then normalised in counts per ten thousand molecules [cptt] throughout. Chromatin accessibility data was binarised.

### Cell type annotation processing

The original cell type annotations were inspected from the data/metadata objects in conjunction with the original publication, especially embeddings. For transcriptomic data, expression of marker genes for specific cell types were verified manually: when markers for a distinct cell type were found instead of the expected ones, the cell type was reannotated based on the markers found together with a literature search on that particular organism and organ. Whenever the original publication used subtyping with no biological meaning (e.g. “fibroblast II”) or no interpretation was given in the publication main text either, the cell type was reannotated at a coarser level (e.g. “fibroblast”), but only if the embedding of distinct subtypes was relatively contiguous - which indicates absence of obvious and potentially biologically meaningful clusters. Cells with type annotations that were obviously labelled for discard (e.g. “low-quality”) were filtered out.

### Lossy compression algorithms

Two approximation algorithms with different compression ratios were developed. In algorithm 1, cptt-normalised counts from all cells within each reannotated cell type were averaged. The fraction of cells with non-zero signal from each feature was also recorded. In algorithm 2, the original cell embeddings were used or regenerated. K-means was performed at decreasing K, starting from 5 times the number of cell types, until no clusters with only a few cells were found. Average and fraction of cells with non-zero signal were computed within each cluster, as were cluster centroids and convex hulls. For both algorithms, storage for our cloud API server was achieved using species-specific HDF5 files with level-22 zStandard compression [35] and chunked storage [33]. For chromatin accessibility data, no fractions were computed, while averages were quantised into 8-bit unsigned integers with uneven binning (almost logarithmic). Quantisation was undone on the fly when data was accessed via the APIs.

### Application programming interfaces (APIs)

Machine access to atlas approximations was built as a REST (representational state transfer) application programming interface (API) over the HTTP protocol. The API accepts GET requests only, with a unique resource identifier (URI) encoded through parametric addresses (URLs). Cross-origin requests are generally accepted, with a limitation of 50 features per request to encourage users to ask specific questions rather than request full database dumps. Compressed HDF5 files containing entire approximations at once are available as well as documented (see main text). The APIs are served by an Amazon Lightsail cloud instance running a Flask backend [42] within a Docker container [43]. Documented API grammar, available at https://atlasapprox.readthedocs.io, is the only way to interact with the server as by REST principles. While REST API are a design bottleneck to ensure consistency, software packages in Python (*atlasapprox*), R (*atlasapprox*), JavaScript (*@fabilab/atlasapprox*) and Bash (https://github.com/fabilab/cell_atlas_approximations_API/blob/main/shell/atlasapprox) have been created as well to simplify (e.g. via caching) access from those programming languages. Responses are provided by the REST API in JSON format, however the language-specific packages modify those responses to create a more cohesive experience with typical constructs of each language (e.g. in Python, pandas dataframes). The JavaScript package can be used from both nodejs and a browser frontend via Babel or any other CommonJS module transpiler.

### Natural language processing (NLP)

The chatbot uses nlp.js [44] to understand user intent and extract entities such as organisms and organs of choice and feature names. Slot filling is used to request for additional information when a user question is incomplete, which gives a natural feeling to the conversation. Once a question intent and required entities match an API call, the required data is accessed via the JavaScript API using ES6 modules. If the answer is short, it is phrased into text and returned by the bot. If the answer is long and complex (e.g. a table of gene expression), the answer is delegated to a third party. Internally, the chatbot uses a neural network architecture that is pre-trained during package version release to minimise loading times on the user browser. The chatbot is available on npm as *@fabilab/atlasapprox-nlp*.

### Web interface

The web interface is engineered on top of React [45]. Plots such as heatmaps, dot plots, and bar plots are built using plotly.js [46]. A state variable inside an abstract React component keeps the chatbot and the plotting section in sync, with a one-directional flow of information from the chat (i.e. user and bot) towards the plot. Changes in plot appearance (e.g. logarithm of the data, addition or exclusion of features) are all managed through the chatbot. Care has been taken to minimise the number of buttons on the interface to reduce the risk of confusion for the user.

## Data and code availability

The umbrella GitHub repository at https://github.com/fabilab/cell_atlas_approximations is the entry point for code explorations and contains links to other repositories for the (i) approximation algorithms and scripts, (ii) APIs, (iii) natural language processing, and (iv) human interface. All code is open source, documented, and reasonably tested. The APIs are documented at https://atlasapprox.readthedocs.io. The original data is available at their various online locations as specified in the data sources (see API call). HDF5 files with the approximations for each organism are available on FigShare at https://figshare.com/account/projects/191094/articles/24932748.

## Acknowledgements

We would like to express our heartfelt thanks to Robert C Jones, Carsten Knutsen, Cristina Alvira, David Cornfield, Emily Wong, Junyue Cao, Bo Wang, Dania Nanes Sarfati, Simon Peters, Jules Duruz, and all authors of the original cell atlas publications for access to the cell atlas data and for scientific discussions. We would also like to thank Danny Dien for early prototypes of the chatbot. This work is supported by a Chan Zuckerberg Initiative Single Cell Data Insight Grant #DI2-0000000091.

## Supplementary Materials

**Supplementary Figure 1:**
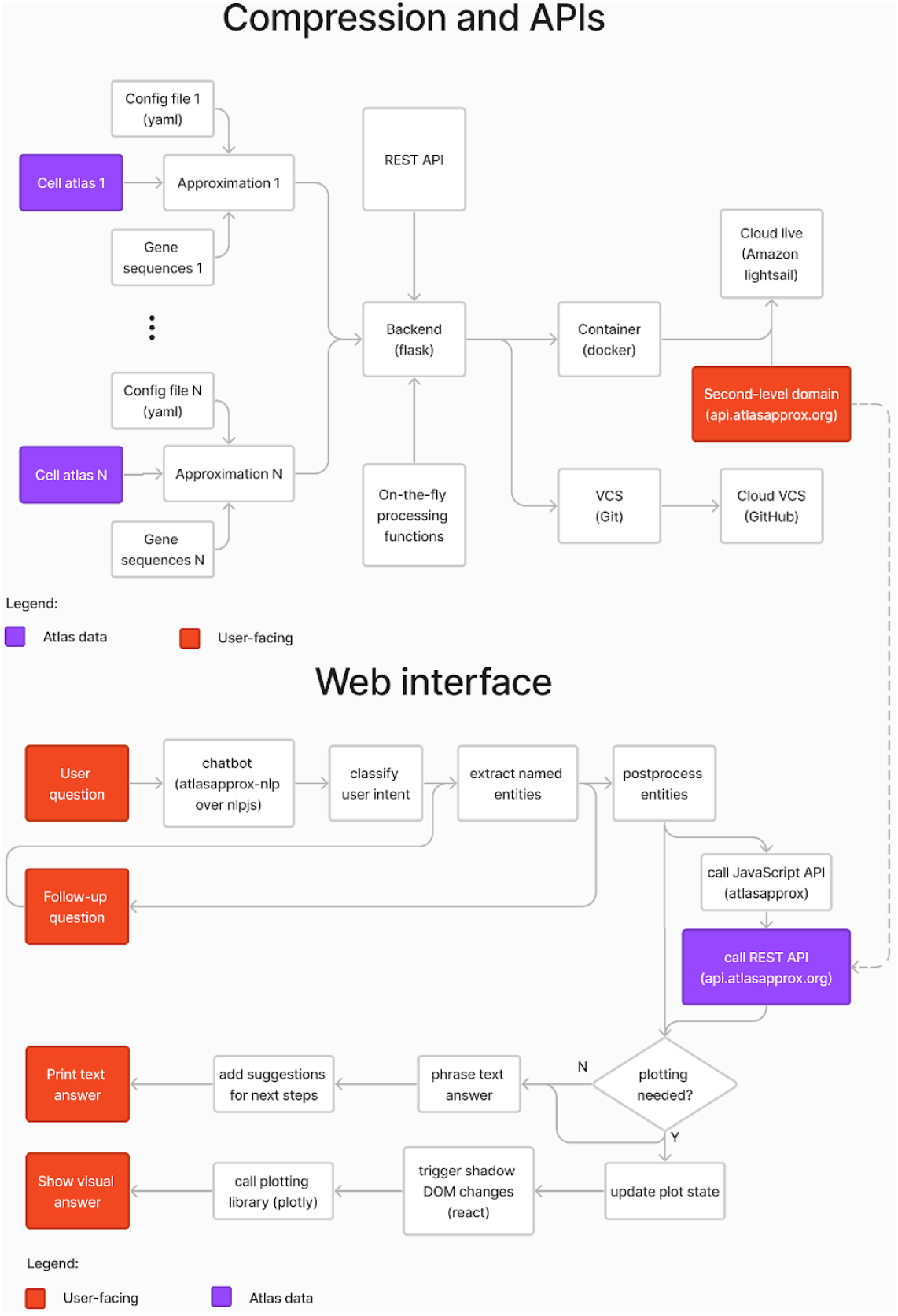
Flow chart of the cell atlas approximations software stack.

## References

1. Quake SR. A decade of molecular cell atlases. Trends Genet. 2022;38: 805–810.

2. Tabula Muris Consortium. Single-cell transcriptomics of 20 mouse organs creates a Tabula Muris. Nature. 2018. doi:10.1038/s41586-018-0590-4

3. Tabula Sapiens Consortium* Jones RC, Karkanias J, Krasnow MA, Pisco AO, Quake SR, et al. The Tabula Sapiens: A multiple-organ, single-cell transcriptomic atlas of humans. Science. 2022;376:eabl4896.

4. Li P, Nanes Sarfati D, Xue Y, Yu X, Tarashansky AJ, Quake SR, et al. Single-cell analysis of Schistosoma mansoni identifies a conserved genetic program controlling germline stem cell fate. NatCommun. 2021;12: 485.

5. Sebé-Pedrós A, Chomsky E, Pang K, Lara-Astiaso D, Gaiti F, Mukamel Z, et al. Early metazoan celltype diversity and the evolution of multicellular gene regulation. Nat Ecol Evol. 2018;2: 1176–1188.

6. Zhang K, Hocker JD, Miller M, Hou X, Chiou J, Poirion OB, et al. A single-cell atlas of chromatin accessibility in the human genome. Cell. 2021;184: 5985–6001.e19.

7. The Tabula Microcebus Consortium, Ezran C, Liu S, Chang S, Ming J, Botvinnik O, et al. Tabula Microcebus: A transcriptomic cell atlas of mouse lemur, an emerging primate model organism. bioRxiv.2022. p. 2021.12.12.469460. doi:10.1101/2021.12.12.469460

8. Cao J, Packer JS, Ramani V, Cusanovich DA, Huynh C, Daza R, et al. Comprehensive single-cell transcriptional profiling of a multicellular organism. Science. 2017;357: 661–667.

9. Wagner DE, Weinreb C, Collins ZM, Briggs JA, Megason SG, Klein AM. Single-cell mapping of geneexpression landscapes and lineage in the zebrafish embryo. Science. 2018;360: 981–987.

10. Musser JM, Schippers KJ, Nickel M, Mizzon G, Kohn AB, Pape C, et al. Profiling cellular diversity in sponges informs animal cell type and nervous system evolution. Science. 2021;374: 717–723.

11. Liao Y, Ma L, Guo Q E W, Fang X, Yang L, et al. Cell landscape of larval and adult Xenopus laevis at single-cell resolution. Nat Commun. 2022;13: 4306.

12. Plass M, Solana J, Wolf FA, Ayoub S, Misios A, Glažar P, et al. Cell type atlas and lineage tree of awhole complex animal by single-cell transcriptomics. Science. 2018;360. doi:10.1126/science.aaq1723

13. McInnes L, Healy J, Melville J. UMAP: Uniform Manifold Approximation and Projection for DimensionReduction. arXiv [stat.ML]. 2018. Available: http://arxiv.org/abs/1802.03426

14. Ouyang JF, Kamaraj US, Cao EY, Rackham OJL. ShinyCell: simple and sharable visualization of single-cell gene expression data. Bioinformatics. 2021;37: 3374–3376.

15. Chan Zuckerberg Initiative. CZ CELLxGENE Discover. In: CZ CELLxGENE Discover [Internet]. [cited 14 Aug 2023]. Available: https://cellxgene.cziscience.com/

16. Chen Y, Zhang X, Peng X, Jin Y, Ding P, Xiao J, et al. SPEED: Single-cell Pan-species atlas in the light of Ecology and Evolution for Development and Diseases. Nucleic Acids Res. 2023;51: D1150–D1159.

17. Ghita L, Yao Z, Xie Y, Duran V, Cagirici HB, Samir J, et al. Global and cell type-specific immunological hallmarks of severe dengue progression identified via a systems immunology approach. Nat Immunol.2023;24: 2150–2163.

18. Yao Z, Zanini F, Kumar S, Karim M, Saul S, Bhalla N, et al. The transcriptional landscape of Venezuelan equine encephalitis virus (TC-83) infection. PLoS Negl Trop Dis. 2021;15: e0009306.

19. Zanini F, Robinson ML, Croote D, Sahoo MK, Sanz AM, Ortiz-Lasso E, et al. Virus-inclusive single-cellRNA sequencing reveals the molecular signature of progression to severe dengue. Proc Natl Acad SciU S A. 2018;115: E12363–E12369.

20. Domingo-Gonzalez R, Zanini F, Che X, Liu M, Jones RC, Swift MA, et al. Diverse homeostatic and immunomodulatory roles of immune cells in the developing mouse lung at single cell resolution. Elife.2020;9. doi:10.7554/eLife.56890

21. Zanini F, Che X, Knutsen C, Liu M, Suresh NE, Domingo-Gonzalez R, et al. Developmental diversity and unique sensitivity to injury of lung endothelial subtypes during postnatal growth. iScience. 2023;26:106097.

22. Thoms JAI, Knezevic K, Harvey G, Huang Y, Seneviratne JA, Carter DR, et al. Disruption of a GATA2,TAL1, ERG regulatory circuit promotes erythroid transition in healthy and leukemic stem cells. ColdSpring Harbor Laboratory. 2020. p. 2020.10.26.353797. doi:10.1101/2020.10.26.353797

23. Tabula Muris Consortium. A single-cell transcriptomic atlas characterizes ageing tissues in the mouse.Nature. 2020;583: 590–595.

24. Wolf FA, Angerer P, Theis FJ. SCANPY: large-scale single-cell gene expression data analysis. Genome Biol. 2018;19: 15.

25. Hao Y, Stuart T, Kowalski MH, Choudhary S, Hoffman P, Hartman A, et al. Dictionary learning for integrative, multimodal and scalable single-cell analysis. Nat Biotechnol. 2023.doi:10.1038/s41587-023-01767-y

26. Harris CR, Millman KJ, van der Walt SJ, Gommers R, Virtanen P, Cournapeau D, et al. Array programming with NumPy. Nature. 2020;585: 357–362.

27. Virtanen P, Gommers R, Oliphant TE, Haberland M, Reddy T, Cournapeau D, et al. SciPy 1.0:fundamental algorithms for scientific computing in Python. Nat Methods. 2020;17: 261–272.

28. Hunter JD. Matplotlib: A 2D graphics environment. Comput Sci Eng. 2007;9: 90–95.

29. McKinney W. pandas: a Foundational Python Library for Data Analysis and Statistics. 2011.

30. Antonov M, Csárdi G, Horvát S, Müller K, Nepusz T, Noom D, et al. igraph enables fast and robust network analysis across programming languages. arXiv [cs.SI]. 2023. Available: http://arxiv.org/abs/2311.10260

31. Granger BE, Pérez F. Jupyter: Thinking and Storytelling With Code and Data. Comput Sci Eng.2021;23: 7–14.

32. Introducing ChatGPT. [cited 28 Dec 2023]. Available: https://openai.com/blog/chatgpt

33. The HDF Group. Hierarchical Data Format, version 5. In: Hierarchical Data Format, version 5 [Internet].1997 [cited 14 Aug 2023]. Available: https://www.hdfgroup.org/HDF5

34. Lin Z, Akin H, Rao R, Hie B, Zhu Z, Lu W, et al. Evolutionary-scale prediction of atomic-level proteinstructure with a language model. Science. 2023;379: 1123–1130.

35. Zstandard. [cited 14 Aug 2023]. Available: http://facebook.github.io/zstd/

36. Dwivedi YK, Kshetri N, Hughes L, Slade EL, Jeyaraj A, Kar AK, et al. Opinion Paper: “So what if ChatGPT wrote it?” Multidisciplinary perspectives on opportunities, challenges and implications of generative conversational AI for research, practice and policy. Int J Inf Manage. 2023;71: 102642.

37. Alkaissi H, McFarlane SI. Artificial Hallucinations in ChatGPT: Implications in Scientific Writing. Cureus.2023;15: e35179.

38. Zanini F, Berghuis BA, Jones RC, Nicolis di Robilant B, Nong RY, Norton JA, et al. Northstar enables automatic classification of known and novel cell types from tumor samples. Sci Rep. 2020;10: 15251.

39. Lin Y, Cao Y, Kim HJ, Salim A, Speed TP, Lin D, et al. scClassify: hierarchical classification of cells.bioRxiv. 2019. p. 776948. doi:10.1101/776948

40. Lopez R, Regier J, Cole MB, Jordan MI, Yosef N. Deep generative modeling for single-cell transcriptomics. Nat Methods. 2018;15: 1053–1058.

41. Redelings BD, Holder MT. A supertree pipeline for summarizing phylogenetic and taxonomic information for millions of species. PeerJ. 2017;5: e3058.

42. Grinberg M. Flask Web Development: Developing Web Applications with Python. 2nd ed. O’Reilly Media; 2018.

43. Merkel D. Docker: lightweight Linux containers for consistent development and deployment. Linux J.2014;2014: 2.

44. nlp.js: An NLP library for building bots, with entity extraction, sentiment analysis, automatic language identify, and so more. Github; Available: https://github.com/axa-group/nlp.js

45. React. [cited 14 Aug 2023]. Available: https://react.dev/

46. Plotly Technologies Inc. Collaborative data science. In: Collaborative data science [Internet]. Montréal,QC: Plotly Technologies Inc.; 2015 [cited 14 Aug 2023]. Available:https://plot.ly

